# A Robust and Scalable Graph Neural Network for Accurate Single Cell Classification

**DOI:** 10.1101/2021.06.24.449752

**Authors:** Yuansong Zeng, Xiang Zhou, Zixiang Pan, Yutong Lu, Yuedong Yang

## Abstract

**Motivation:** Single-cell RNA sequencing (scRNA-seq) techniques provide high-resolution data on cellular heterogeneity in diverse tissues, and a critical step for the data analysis is cell type identification. Traditional methods usually cluster the cells and manually identify cell clusters through marker genes, which is time-consuming and subjective. With the launch of several large-scale single-cell projects, millions of sequenced cells have been annotated and it is promising to transfer labels from the annotated datasets to newly generated datasets. One powerful way for the transferring is to learn cell relations through the graph neural network (GNN), while vanilla GNN is difficult to process millions of cells due to the expensive costs of the message-passing procedure at each training epoch.

**Results:** Here, we have developed a robust and scalable GNN-based method for accurate single cell classification (GraphCS), where the graph is constructed to connect similar cells within and between labelled and unlabelled scRNA-seq datasets for propagation of shared information. To overcome the slow information propagation of GNN at each training epoch, the diffused information is pre-calculated via the approximate Generalized PageRank algorithm, enabling sublinear complexity for a high speed and scalability on millions of cells. Compared with existing methods, GraphCS demonstrates better performance on simulated, cross-platform, and cross-species scRNA-seq datasets. More importantly, our model can achieve superior performance on a large dataset with one million cells within 50 minutes.

## 1. Introduction

Single cell RNA-seq technologies promise to provide high-resolution insights into the complex cellular ecosystem[1–3] by measuring gene expression in millions of single cells from multiple samples [4–8]. Several large-scale single-cell projects, e.g. the Human Cell Atlas, have been established as a result of the decreasing costs in scRNA-seq technologies [9, 10]. In scRNA-seq studies, an essential step is to identify the sequenced cells through the sequenced gene expression [11], which is usually obtained through cell clustering and subsequently manually identifying cell clusters through marker genes[12]. This process is time-consuming and subjective.

With the tremendous increase of well-annotated scRNA-seq datasets, it is feasible to transfer well-defined labels (cell types) of existing single-cell datasets to newly generated single-cell datasets [13, 14]. However, the knowledge transferring is challenging due to various noises among scRNA-seq data (e.g., dropout) [15, 16]. In addition, batch effects exist between single-cell datasets because they are usually collected from different platforms[17, 18], tissues, or species[19, 20]. Early methods were developed to search for similar cells in the reference datasets with well-defined labels. For example, scmap[21] measures the maximum similarity between well-annotated cells of reference data and unknown query data to annotate cell types. SingleR[22] measures the similarity by calculating the correlation between gene expression. CHETAH[23] identifies the unknown cells using the high cumulative density of each cell type correlation distribution. Obviously, these methods consider only pairwise similarity and have ignored the non-linear relations between annotated cells. For this reason, several methods train classifiers using the labeled datasets or reference atlas, and make predictions on query datasets. For example, scPred [24] trains a support vector machine by using the features obtained from singular value decomposition. SingleCellNet [25] applies an ensemble of boosted regression trees and a Random Forest classifier to annotate cells. Seurat[26] is a commonly used toolkit in single-cell studies, which applies a specialized method to transfer labels to unknown cell types. Though valuable in different scenes, these methods still exhibit limited performance, partially due to their ignoration of higher-order relations between cells.

In fact, the high-order representation and topological relations could been naturally learned by the graph neural network (GNN), and GNN have been proven with improved performance in scRNA-seq data analyses such as imputation and clustering [27–29]. ScGCN[30] is currently the only graph neural network method for annotating cells. The method is based on the GNN architecture proposed by Kipf and Welling [31], which relies on an expensive message-passing procedure to propagate information and has to include the full-batch during training. Thus, the huge costs of computations and memory prevent its applications to large datasets, especially with the arrival of datasets containing millions of cells [32, 33].

To solve the scalability of GNN, many studies have been proposed. For example, Chen et al. [34] proposed a scalable GNN model, which could be efficiently trained with mini-batches using GPU. One critical point is its approximation of the diffused information through the bidirectional propagation by the Generalized PageRank algorithm [35], which avoids iterative information diffusion in each training epoch. In addition, the use of mini-batch training reduces the requirement of large GPU memory from full-batch training. Thus, the method could be used on large graphs with billions of edges.

Another issue for GNN is to accurately construct the cell graph among millions of cells. Traditional methods such as Cosine similarity, KNN, UMAP [36], and Annoy [37] (https://github.com/spotify/annoy) are widely used for constructing the cell graph by measuring the cell-to-cell similarity in single-cell RNA-seq data [38–40], but they do not take account of the batch effects between datasets. To consider the batch effects, several methods captures the cell relations through scGCN [30] constructs the cell graph using CCA-MNN, a combination of canonical correlation analysis (CCA) [41] and the mutual nearest neighbour (MNN) [42]. Conos [43] relies on multiple plausible inter-sample mappings to construct a graph connecting all measured cells. BBKNN [44] provides an extremely fast and scalable neighbourhood construction method across all batches. The runtimes of BBKNN scale linearly with the increase in number of cells through integrating the approximate neighbor detection technique in algorithm Annoy.

Here, we present a scalable graph neural network learning model for cell annotations by constructing the graph via BBKNN, and pre-calculate the diffused features via the graph bidirectional propagation algorithm (GBP). Concretely, GBP propagates information among similar cells within and between labeled and unlabeled datasets, resulting in significant gains of speed and scalability of GNN while efficiently removing the batch effects. The integrated features from the GBP module are then inputted to a classification neural network to annotate cells for the query dataset. To better estimate the decision boundary between different cell types, we also use the virtual adversarial training (VAT) loss [45] to improve model generality. Our method was demonstrated to outperform other methods on both simulated datasets and real datasets across species and platforms. More importantly, the model can be extended to large-scale datasets in a reasonable time scale.

## 2. Materials and Methods

### 2.1 Datasets and Pre-processing

#### Simulated datasets

We simulated different batches of scRNA-seq data through the R package “Splatter” [46]. Specifically, we used Splatter to generate paired batches as the reference and query dataset. Each simulated dataset was comprised of four cell groups with the same cell proportion by setting the parameter group.prob of value 0.25. We fixed the number of cells in the reference and query data respectively as 2000 and 1000 both with the number of genes as 10000. To simulate different magnitudes of batch effects in scRNA-seq data, we tested different batch.facScale values from {0.2, 0.4, 0.6, 0.8,1.0,1.2,1.4, 1.6}, where greater values of batch.facScale correspond to larger batch effects between the reference and query datasets. For other parameters, default values were applied unless otherwise specified. We generated five simulated datasets from different random seeds and reported the average results.

#### Cross-species datasets

The cross-species datasets consisted of four paired cross-species pancreas datasets downloaded from the SingleCellNet GitHub page. Specifically, we used the mouse pancreas dataset generated by Baron et al[1] as the reference data and the human pancreas datasets generated by Baron et al[1] and Segerstolpe et al[47] as the query data, respectively. We also used the Baron mouse pancreas dataset to annotate a combination dataset containing five human pancreas datasets respectively generated by Baron et al, Wang et al[48], Xin et al[49], Muraro et al[6], and Segerstolpe et al. Finally, we used the human pancreas dataset generated by Baron et al as the reference data and the mouse pancreas dataset also generated by Baron et al as the query data. The compatible genes between species were obtained through homologous genes convert interface provided by SingleCellNet[25].

#### Multiple reference datasets

To investigate the performance of our model on multiple reference datasets, we integrated the human pancreas datasets by Wang et al, Muraro et al, and Segerstolpe et al as the reference to annotate the Baron human pancreas dataset. Specifically, we removed the acinar and alpha cells from datasets by Segerstolpe et al and Wang et al, only reserving them in the Muraro et al dataset. We also removed the endothelial cells from Muraro et al, only reserving them in the Segerstolpe et al dataset. Thus, each reference dataset has its own unique cell type, while the query dataset Baron et al includes all these unique cell types: cell acinar, alpha, and endothelial.

#### Cross-platform datasets

We used seven paired cross-platform datasets in this study. The first one is the Peripheral Blood Mononuclear Cells (PBMC) scRNA-seq data from the SeuratData package with dataset name “pbmcsca” [50], which is widely used for evaluating cell annotating methods. PBMC consisted of seven batches from seven different sequencing platforms: 10x Chromium (v2), 10x Chromium (v3), Seq-Well, Smart-seq2, inDrop, Drop-seq, and CEL-Seq2. Two 10x Chromium datasets were combined as the reference data, and the rest five datasets were used as the query data. Thus, PBMC has five paired cross-platform datasets. Then, in order to evaluate the scalability of our method, we downloaded two large cross-platform datasets: the mouse retina and mouse brain datasets from ref [51]. Specifically, the mouse retina dataset contained two batches that were generated based on the Drop-seq technology by two unassociated laboratories [52, 53]. These two mouse retina datasets contained 26,830 and 44,808 cells, and were used as the reference and query data, respectively. The mouse brain data contained two batches that were generated by the Drop-seq and SPLiT-seq protocols, respectively [39]. They were used as the reference and the query data that contained 691,600 and 141,606 cells, respectively.

#### Preprocessing

All simulated datasets were normalized through the transcripts per million (TPM) method [54]. For real datasets, we followed the standard procedure proposed in Seurat to normalize the gene expression matrix. Specifically, the “NormalizeData” function was run with the default parameter “LogNormalize” and the scaling factor of 10,000. Then, we selected the top 2000 highly variable genes based on the normalized matrix through the “FindVariableFeatures” function.

### 2.2 The architecture of GraphCS

This study proposed a robust and scalable graph neural network model to annotate cell types in a semi-supervised manner. As shown in Fig. 1, the GraphCS model consists of three modules: graph construction, graph bi-directional propagation (GBP), and classification modules.

**Fig. 1.**
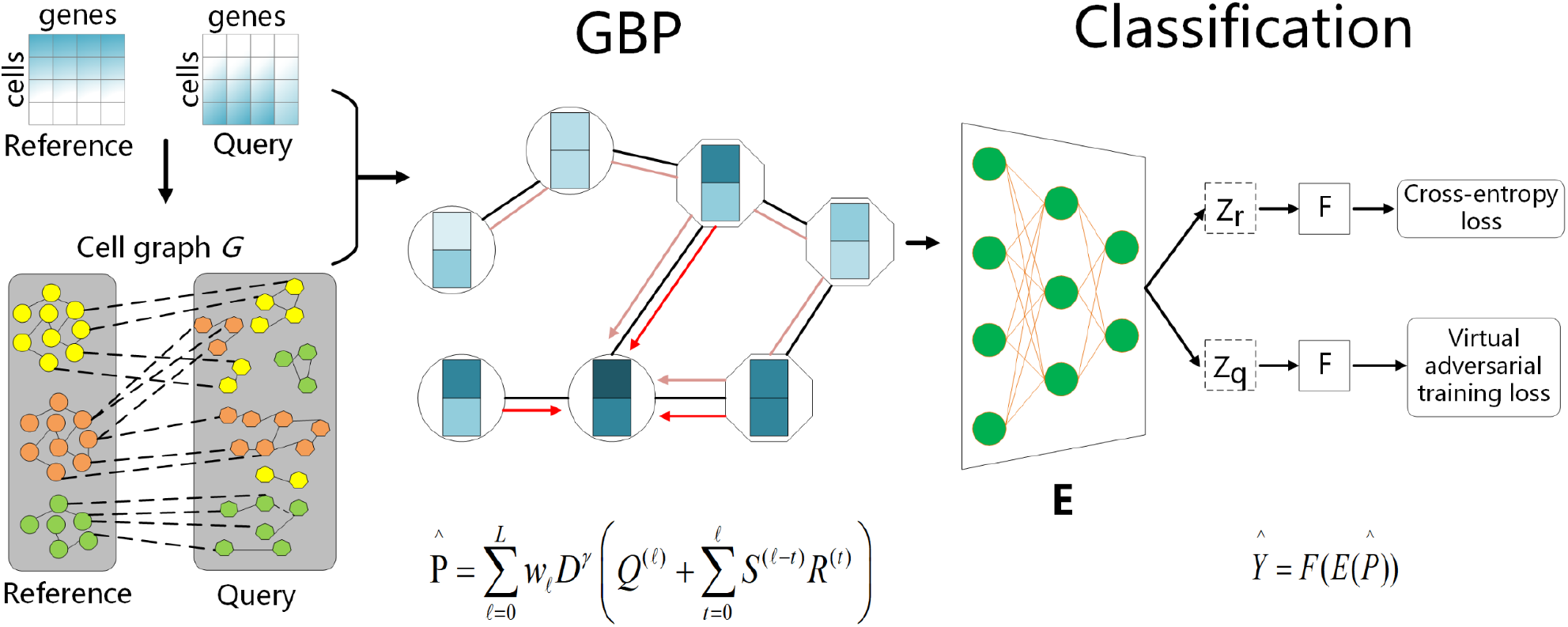
The Schematic overview of GraphCS for cell type classification. GraphCS consists of graph construction, graph bidirectional propagation (GBP), and classification modules. The graph construction module constructs the cell graph according to gene expression similarity through the BBKNN algorithm. Through the graph, the GBP module diffuses feature information among cells, which is then inputted into the classification module used to classify cells.

#### 2.2.1 Graph construction module

The cell graph *G* is constructed by linking cells with similar gene expressions within and between the reference and query datasets. Here, we construct the graph *G* by BBKNN with default parameters, which provides a fast and scalable neighbourhood construction method across all batches. Briefly, for each cell *c*, three most similar cells are selected with the lowest Euclidean distances from each of *N_b_* batches (including the batch itself). The connected cell graph is then inputted into UMAP for recalculating connectivity scores, through which neighboured cells are trimmed so that each cell contains at most 30*N_b_* neighbours (edges).

#### 2.2.2 GBP module

To acquire high scalability, GNN is estimated through the Generalized PageRank algorithm, which is further approximated by the Graph Bidirectional Propagation Algorithm.

##### Generalized PageRank Algorithm

To acquire high scalability of GNN, the feature propagation is pre-calculated through Generalized PageRank matrix as:

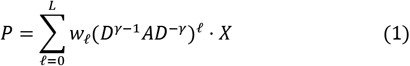

where w_*ℓ*_ is the weight of the *ℓ*-th order convolution matrix, *A* and *D* are the adjacency matrix and diagonal degree matrix of graph *G*, respectively, *X* is the feature matrix, and *γ* is the convolution coefficient. This strategy has been proven to well estimate feature propagation [55], and we followed the study to set *w*_*ℓ*_ = α(1 − α)^*ℓ*^ for constant decay factor α ∈ (0,1).

##### The Graph Bidirectional Propagation Algorithm

To reduce the time complexity, the Generalized PageRank is further approximated with the graph bidirectional propagation that combines the Monte-Carlo Propagation and Reverse Push Propagation. Graph bidirectional propagation has been proven to provide accurate unbiased estimator [34]. Concretely, we use the following formulate as an unbiased estimator for the Generalized PageRank matrix *P* defined in equation (1).

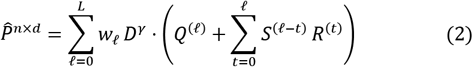

where *n* is the total number of cells in the reference and the query data, and *d* is the size of gene features. *Q* and *R* are respectively the reserve matrix and the residue matrix originated from the Reverse Push Propagation algorithm, and *S* records the fraction of random walks from the Monte-Carlo Propagation. The detailed information and proof of Equation (2) can be found in ref [34].

#### 2.2.3 Classification Module

After obtaining the feature matrix 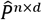 from the GBP module, we apply a neural network classification with mini-batch training to make the cell type prediction. In addition, the virtual adversarial training is employed to improve the generality of our model.

##### Neural Network Classification

Our classification module contains a neural network feature extractor *E* with multiple hidden layers and a label predictor F with a *Softmax* output layer. The input of classification module includes reference gene expression matrix 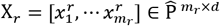 with the corresponding labels 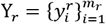 and query gene expression matrix 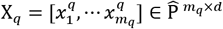. We optimize the classification module using the following standard cross-entropy loss:

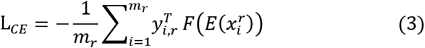

where y_*i,r*_ ∈ R^*CL*×1^ is one-hot encoded vector of 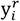 *CL* is the number of class.

##### Virtual Adversarial Training

Virtual adversarial training (VAT) is applied to improve the generalization of the classification module by incorporating the information of data distribution from query data. VAT is a data augmentation technique without prior label information[56]. VAT tries to make predictions invariant to small perturbation by minimizing the distance between the input and a perturbed version of the input. Then the model is robust to small noises or perturbations in the inputs. We compute VAT’s loss function as the following:

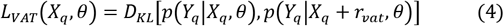

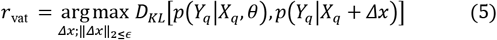

where *r_vat_* optimizes the difference between the model output of the non-perturbed input and the perturbed input, *θ* is parameter of the model, *Δx* is a Gaussian noise and *Y_q_* is predicted by the label predictor F. The hyper-parameter *ϵ* is the norm constraint for the adversarial direction, and we set *ϵ* to 0.1 following the previous study [45]. The output distribution is parameterized as *p*(Y_*q*_|X_*q*_, *θ*), and D_*KL*_ [∙,∙] is KullbackLeibler divergence.

So, the total loss function of classification module as the following:

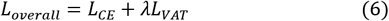

where λ is the hyper-parameters to balance the contribution of virtual adversarial training to the total loss function.

### 2.3 Model interpretation by selecting key features

In order to obtain important genes for each predicted cell type, we estimate gene importance based on the gradient of the correct category logit with respect to the input vector used the activation maximization method [57, 58]. Specifically, given the trained neural network φ and a predicted cell type *i*, activation maximization searches for key input genes *x*^∗^ by solving the following optimization problem:

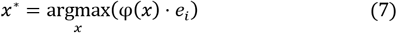

where *e_i_* is the *i*-th category’s natural basis vector. The formulation of (7) can be solved by backpropagation, where the gradient of φ(*x*) with respect to *x* are computed to update the input *x* iteratively. Specifically, we initialize the optimization with a zero vector and then run the optimization for 100 iterations with a learning rate of 1 as recommended in ref [59]. Those inputs leading to the largest changes in comparison with the initialization values are selected as the important genes. To evaluate the identified top-important genes, we conduct the GO analysis on the selected top genes with the largest changes using the R package clusterProfiler[60].

### 2.4 Hyper-parameters setting

The GraphCS was implemented in PyTorch and C++. For GBP module, we set *α*=0.05 and *γ*=0.5 for all datasets. For classification module, we set the number of neural network layers and the learning rate as 2 and 0.001, respectively. For the contribution of VAT loss, we set λ=0.1 for all datasets. We set the training batch size of classification module as 1024 and 4096 when the total number of cells exceeds 10000 and 50000, respectively; In other situations, the training batch size is set as 128. For cross-species datasets, the maximum of inter edges in the graph *G* (edges between reference and query data) for each cell is less than four. All results reported in this paper were conducted on Centos 7.0 with Intel® Core (TM) i7-8700K CPU @ 3.70 GHz and 256 GB memory, with the Nvidia Tesla P100 (16G).

### 2.5 Benchmarking classification methods

To evaluate the performance, we compared GraphCS with other tools including: Seurat V3, scmap, scPred, CHETAH, SingleR, SingleCellNet and scGCN. For Seurat V3, we applied both the PCA-based and CCA-based version to evaluate whether the aligned data was benefit for classification. We used the default hyper-parameters recommended in the origin paper for the competing methods.

#### Evaluation metrics

We evaluated the classification performance for all methods using the accuracy, the proportion of correctly annotated cells. For each dataset, we considered the cell type annotations provided by the original dataset as the ground truth.

## 3. Results

### 3.1 Performance on simulated datasets

To investigate the performance of GraphCS under different magnitudes of batch effects, we generated the simulated scRNA-seq data by setting different values of “batch.facScale” through the R package “Splatter”. As shown in Fig. 2a, the accuracies of all methods decreased with the increase of batch.facScale since higher batch.facScale represented larger batch effects, i.e., higher annotating difficulty. Overall, our method consistently achieved stable and the best performance with the accuracies only slightly changed from 1.0 to 0.98 when increasing batch.facScale from 0.2 to 1.6. By comparison, scGCN, the second-best method, had significant drop in accuracies when batch.facScale was greater than 1.0, and a sharp drop from 0.93 to 0.76 when increasing batch.facScale from 1.4 to 1.6. The accuracies of SingleR and scPred were larger than 0.9 when the value of batch.facScale was less than 0.4, but their accuracies significantly dropped afterwards and were only 0.6 and 0.4, respectively when batch.facScale=1.6. SingleCellNet, CHETAH, and scmap, could achieve decent results at batch.facScale of 0.2 with accuracies of 0.87, 0.70, and 0.68, respectively, but they performed badly at batch.facScale of 1.2 with accuracies below 0.5. For two Seurat methods, Seurat-PCA was more sensitive from the batch.facScale value. Seurat-PCA had higher accuracies than Seurat-CCA at batch.facScale of <0.6, but lower accuracies at greater batch.facScale values. This is likely because Seurat-CCA overcorrected the batch effects at small batch.facScale values. By comparison, GraphCS always outperformed the competing methods in different magnitudes of batch effects. The superior performance showed that GraphCS could effectively reduce performance degradation brought by batch difference.

**Fig. 2.**
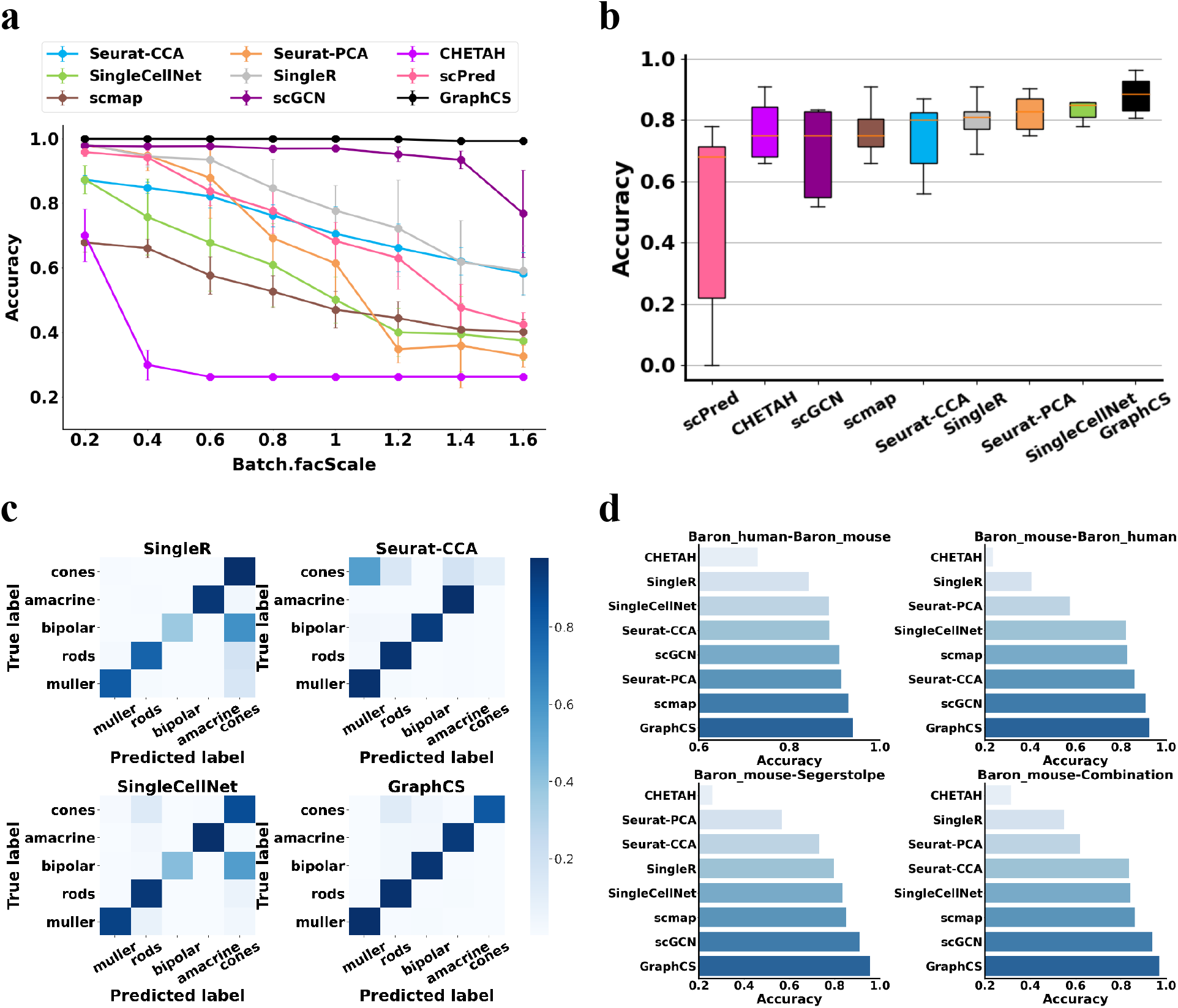
The performance of GraphCS on simulated, cross-platform, and cross-species datasets: **(a)** the average and mean square error values of cell type prediction accuracy on 5 groups of simulated scRNA-seq data at different batch.facScale values; **(b)** the boxplots of cell type prediction accuracy of all methods based on the cross-platform datasets; **(c)** the accuracy matrix of each cell type identified by different methods on the mouse retina dataset; **(d)** the performance of GraphCS on four paired cross-species datasets. Baron_human-Baron_mouse represents the Baron human pancreas dataset as the reference to annotate the Baron mouse pancreas dataset. The rest results represent that the Baron mouse pancreas dataset as the reference to annotate respectively the Baron, Segerstolpe, and the combination human pancreas datasets (the combination contained five human pancreas datasets, including Baron et al, Wang et al, Xin et al, Muraro et al, and Segerstolpe et al). Each bar represents the accuracy of each method.

### 3.2 Performance on real datasets

We evaluated GraphCS on real datasets at two different levels: cross-platform and cross-species. For the cross-platform datasets, we tested on seven paired cross-platform datasets. As shown in Fig. 2b, the average accuracy of GraphCS (mean Acc=88%) was 4% higher than the second-ranked method SingleCellNet (mean Acc=84%) and consistently outperformed other competing methods. Seurat-PCA, SingleR, and Seurat-CCA ranked the 3rd, 4-th, and 5-th in terms of the average accuracy, respectively. In comparison to Seurat-PCA, Seurat-CCA did not benefit from aligning and integrating the datasets. CHETAH, scmap, and scGCN achieved similar average accuracy. Though scGCN took a similar technique to ours, the average accuracy of scGCN was lower than GraphCS. It is likely because the scGCN constructed graphs containing fewer edges (averagely one to two times lower than ours) and didn’t fully utilized the advantages of graph neural network. ScPred achieved much lower performance than other methods. To highlight the comparison regarding specific cell types, we used the heatmap to show the accuracy of each cell type annotated by different methods on the mouse retina dataset. As shown in Fig. 2c and Figure S1. SingleCellNet, SingleR, and scmap incorrectly assigned most of bipolar cells. Seurat-CCA, Seurat-PCA, and CHETAH incorrectly assigned most of cones cells. In contrast, our method correctly discriminated most cell types. Additionally, we performed an experiment by using multiple reference datasets (Figure S2), and our model was shown to consistently outperform other methods.

For the cross-species datasets. We evaluated all methods on four paired cross-species datasets. We didn’t include scPred since it raised exceptions on cross-species datasets. As shown in Fig. 2d, GraphCS achieved an average accuracy of 0.94, respectively 3% and 10% higher than those by the second-ranked method scGCN (0.91) and the third-ranked method scmap (0.84). The left methods are ordered as: SingleCellNet, Seurat-CCA, SingleR, Seruat-PCA, and CHETAH. Specifically, in the combination dataset with only seven T cell and 13 Schwann types, GraphCS could still annotate them accurately (Figure S3a). As shown in the Sankey diagram (Figure S3b), the much smaller number of cells in the reference data than the query data suggests the capability of our model in small reference data.

Finally, we visualized the cells in the aggregated reference-query data of cross-species in Fig. 3. We compared Seurat-CCA, scGCN, and GraphCS since they provided the aggregated data and took account of batch effects between datasets. Cells in the raw data were not separated well due to the substantial noise and batch effects. For example, in the dataset Baron (mouse)-Baron (human), beta cells were separated into two clusters, while alpha and delta cells gathered together. While Seurat and scGCN could discriminate most of the cell populations on all cross-species datasets, they couldn’t explicitly distinguish a few cell types, such as beta and delta cells. By comparison, GraphCS could clearly separate most of the cell populations in all scenarios, indicating its ability to deal with strong batch effects between species.

**Fig. 3.**
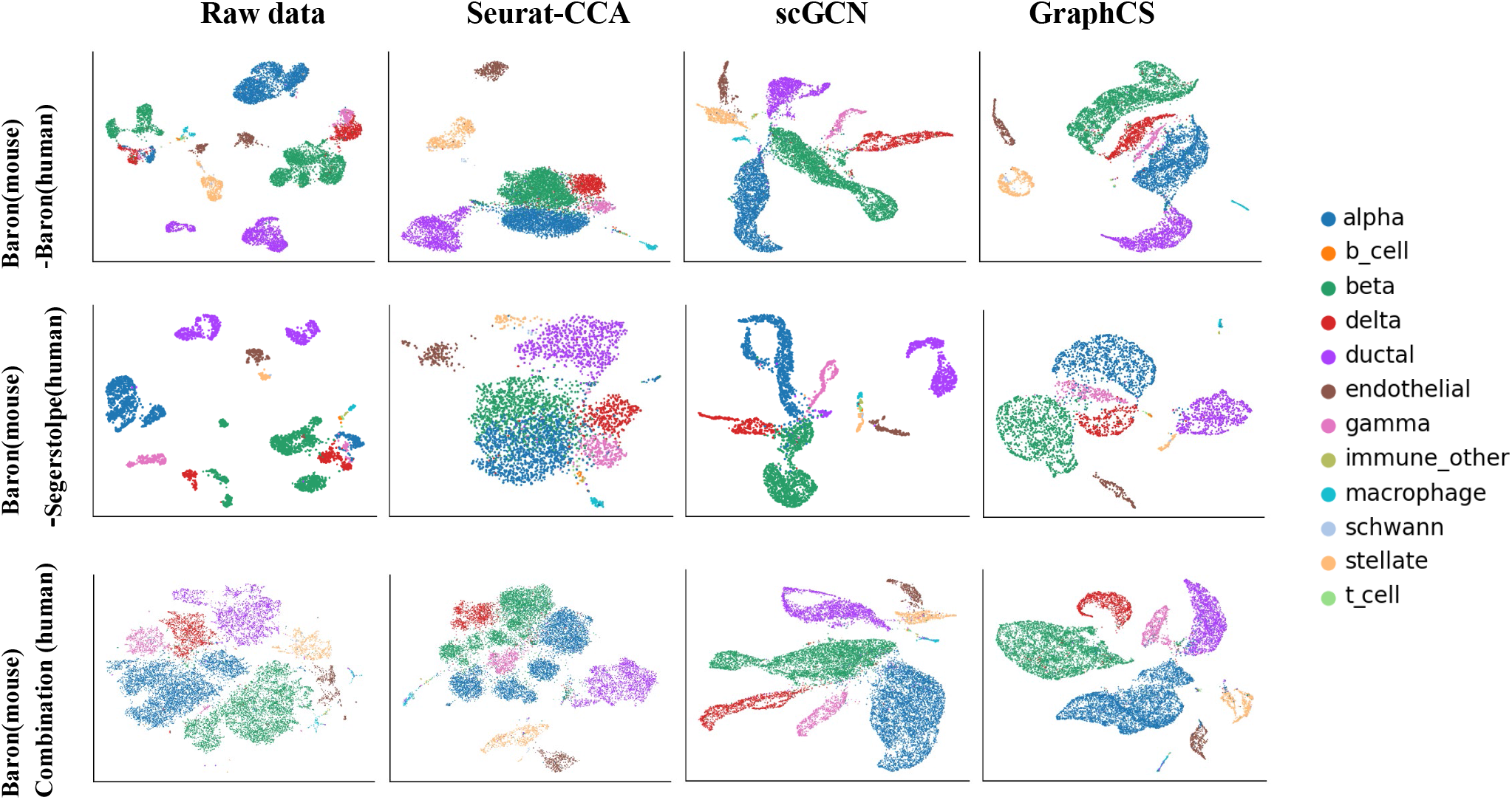
UMAP visualization of three paired cross-species datasets, based on the aggregated data by different methods. The Baron mouse pancreas dataset is the reference for all query datasets. First row: the Baron human pancreas dataset as the query data. Second row: the Segerstolpe human pancreas dataset as the query data. Third row: we combined datasets Baron et al, Wang et al, Xin et al, Muraro et al, and Segerstolpe et al as the query data.

### 3.3 Ablation Experiments

To investigate the contribution of each component in GraphCS, we performed the ablation experiments on cross-species datasets. As shown in Fig. 4, the removal of the GBP module caused a dramatic drop of 18% in the average accuracy, indicating that GBP module efficiently removed the batch effects of inter-datasets by propagating information among similar cells in the graph. The removal of VAT module caused small but significant drop (3.7%) in the average accuracy. In summary, the cooperation of the modules enabled a better annotating of the scRNA-seq data. The trend was similar on the cross-platform datasets (Figure S4).

**Fig. 4.**
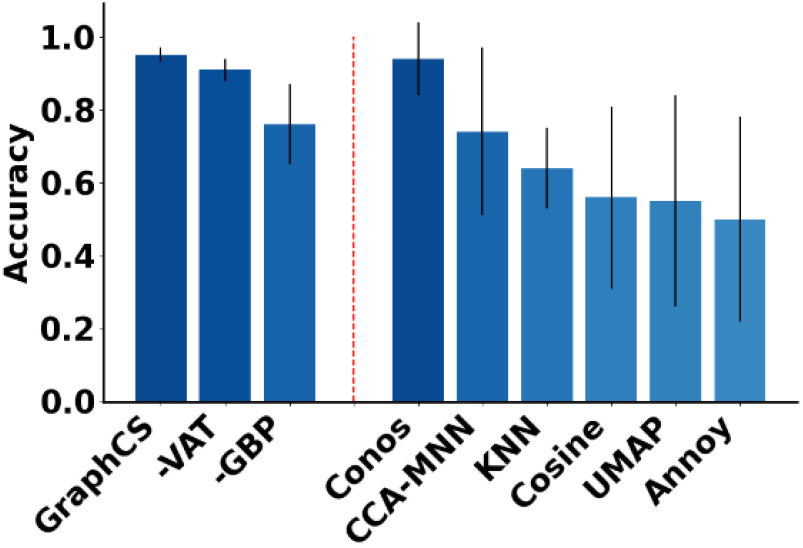
The average classification accuracies and the standard deviations of GraphCS on the cross-species datasets by excluding VAT and GBP modules, and constructing the cell graph by different methods.

We further evaluated the performance of GraphCS using seven different graph construction methods. As shown in Fig. 4, three methods with removal of batch effects (CCA-MNN, Conos, and our method to use BBKNN) achieved higher accuracies than those without removal (Cosine, UMAP, Annoy, and KNN) on the cross-species datasets. BBKNN and Conos achieved similar but better performance than CCA-MNN. It should be noted BBKNN was one to two orders of magnitude faster than Conos and CCA-MNN. The trend was similar on the cross-platform datasets (Figure S5).

### 3.4 Identifying cell-type important genes

To interpret our model, we selected 10 most important genes for each predicted cell type according to the activation maximization method, and used these genes to decide the most enriched GO terms of biological processes through clusterProfiler. As seen in Fig. 5, when using the top 10 genes selected on the cross-platform dataset (10x_Drop-seq), most cell types were significantly enriched on their relevant Go terms. For the B cells, 8 out of top 10 genes (MS4A1, CD79A, BANK1, HLA-DQA1, BLK, CD79B, POU2AF1, and IGHM) were included as maker gene in the PanglaoDB database [61]. These genes were enriched in the Go term “B cell receptor signaling pathway”, consistent with the previous report that the B cell receptor signaling is essential for B cell survival and development although varied in different subpopulations and developmental stages [62]. We also listed top 10 genes for other predicted cell types in Table S1. These results suggested that the identified important genes were consistent with prior knowledge, demonstrating the reliability and interpretability of our model.

**Fig. 5.**
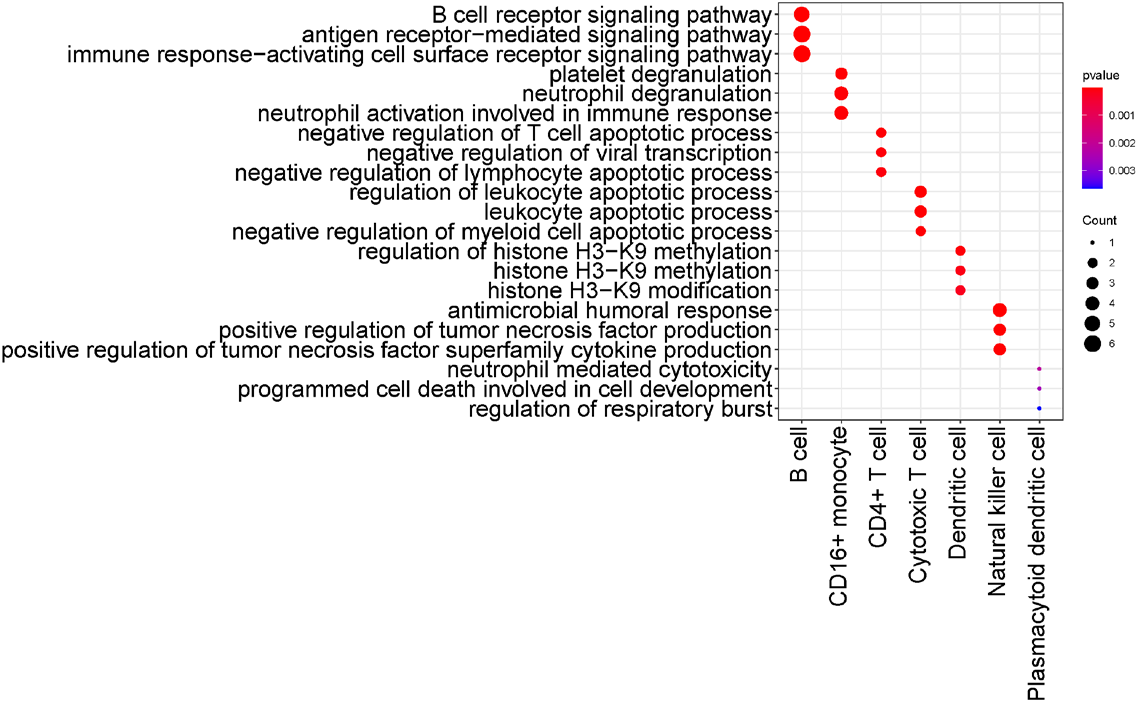
The Enrichment pathways using the top 10 marker genes for each predicted cell type on the 10x_Drop-seq dataset.

### 3.5 Running time evaluation

To evaluate the runtimes of all methods and their scalability with the increase in the number of cells, we sampled the mouse brain dataset in a stratified way (i.e., preserving population frequencies) to 6%, 12%, 36%, 60%, 96%, and 120% of the original number of 833,206 cells and selected the top 2000 highly variable genes as the input features. As shown in Fig. 6, dramatic differences of runtimes could be observed between these methods with increases in the number of cells. GraphCS was faster than all other methods except scmap. GraphCS showed a high scalability with about linear growth of runtimes with the number of cells: 1008s for 500K cells and 2669s for 1000K cells. This was 6 times faster than CHETAH, the next fastest method. Seurate-PCA was close to our method in speed for dataset with 50K cells, but the runtimes dramatically increased when the number of cells with 1000K, and turned 42 times slower than our method. Seurat-CCA was consistently slower than Seurat-PCA, and SingleCellNet was the slowest. While GraphCS was two times slower than scmap, GraphCS consistently achieved average accuracies of 20% higher than scmap. Additionally, under the default parameters, scmap couldn’t process the dataset with >800K cells. Since scGCN constructed the graph based on CCA-MNN that only supported small datasets, it can only run on the small dataset with 50K cells and was 10 times slower than GraphCS. The results demonstrated that our model could be extended to large-scale datasets in linear time complexity.

**Fig. 6.**
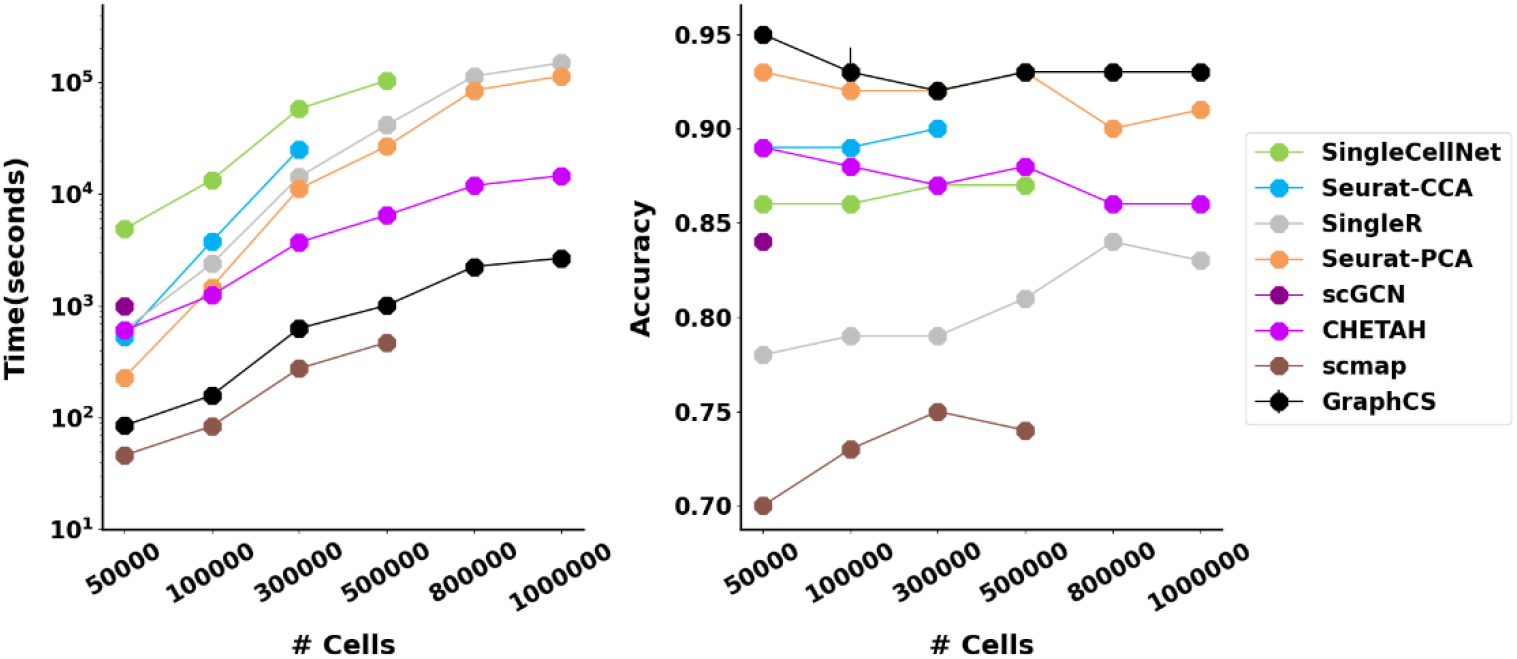
Comparison of different methods for the running time (left) and cell-type classification accuracy (right) on variably sized datasets.

## 4. Discussion

With the tremendous increase of scRNA-seq datasets, it is feasible to transfer well-defined labels of existing single-cell datasets to newly generated single-cell datasets. In this study, we proposed a robust and scalable graph-based artificial intelligence model, which enables training the well-labeled single-cell data to annotate new data through robust knowledge transferring. We have demonstrated that GraphCS achieves significant improvement compared to existing annotation methods in terms of performance and efficiency using the simulated, cross-platform and cross-species scRNA-seq datasets. Meanwhile, our model can be extended to large data in linear time complexity.

While several commonly used cell annotation algorithms, such as Seurat and SingleR, also possess knowledge transferring functionalities, our model achieved superior results in terms of both performance and efficiency. From a technical perspective, our model provides three major advantages. First, GraphCS removes the batch effects between datasets by propagating shared information among neighboring cells. Second, the generality of our model is improved by virtual adversarial training loss by incorporating data distribution information from unlabeled data. Third, GraphCS precomputes the feature propagation of graph neural network in a localized fashion by the graph bidirectional propagation algorithm to achieve scalability.

Although the superior results, GraphCS can be improved in several aspects. Firstly, our model ignores the relations between genes, which has been shown to improve the imputation of scRNA-seq data [27]. Secondly, the performance of our model is influenced by the constructed cell graph, and a high-quality graph can improve performance. Thus, the model may be useful for spatial transcriptomic data analysis [63, 64], where cells could be naturally connected through the provided spatial coordinates. Thirdly, our model highly relies on a high-quality reference dataset: we can’t correctly identify cell types missing in single reference dataset. This problem might be solved by integrating a large number of well-labeled reference datasets since the model was proven able to identify cells simultaneously from two or more reference datasets.

## AVAILABILITY

All source code used in our experiments have been deposited at https://github.com/biomed-AI/GraphCS.

